# Immune, endothelial and neuronal network map in human lymph node and spleen

**DOI:** 10.1101/2021.10.20.465151

**Authors:** Seth Currlin, Harry S. Nick, Marda Jorgensen, Jerelyn A. Nick, Maigan A. Brusko, Hunter Hakimian, Jesus Penaloza-Aponte, Natalie Rodriguez, Miguel Medina-Serpas, Mingder Yang, Robert P. Seifert, Irina Kusmartseva, Todd M. Brusko, Kevin Otto, Amanda L. Posgai, Clive H. Wasserfall, Mark A. Atkinson

## Abstract

The spleen and lymph node represent important hubs for both innate and adaptive immunity^1,2^. Herein, we map immune, endothelial, and neuronal cell networks within these tissues from “normal”/non-diseased organ donors, collected through the NIH Human BioMolecular Atlas Program (HuBMAP)^3^, using highly multiplexed CODEX (CO-Detection by indEXing) imaging and 3D light sheet microscopy of cleared tissues. Building on prior reports^4–6^, we observed the lymph node subcapsular sinus expressing podoplanin, smooth muscle actin, and LYVE1. In the spleen, LYVE1 was expressed by littoral cells lining venous sinusoids, whereas podoplanin was restricted to arteries and trabeculae. 3D visualization of perivascular innervation revealed a subset of axonal processes expressing choline acetyl transferase in both tissues, in contrast with prior literature on human spleen^7^. We further report our novel observations regarding the distinct localization of GAP43 and β3-tubulin within the vascular anatomy of both lymph node and spleen, with Coronin-1A+ cells forming a dense cluster around β3-tubulin positive GAP43 low/negative segments of large vessels in spleen. These data provide an unprecedented 2D and 3D visualization of cellular networks within secondary lymphoid tissues, laying the groundwork for future disease-specific and system-wide studies of neural regulation of immunity in human lymphatics.

## Main

With similarities to the circulation, the lymphatic system stretches across the entire human body^8,9^ to provide the critical functions of conducting lymphatic fluid, filtering blood, mounting defenses against infections and cancer, and shaping immune cell responses to protect against pathogens and autoimmune diseases^8,9^. Tissue organization and function are directly linked in spleen (SPL; blood filtration) and lymph nodes (LN; interstitial fluid filtration) facilitating immune cell priming and activation within follicles as well as afferent and efferent migration through vessels, ducts, and sinusoids^9^. The autonomic nervous system provides control of host immune defense mechanisms through innervation from sympathetic postganglionic neurons, specifically the splenic nerve for the SPL and anatomically paired nerve fibers for respective LNs^10^. However, much of our knowledge regarding the architecture and function of the lymphatic system derives from rodents^11^, with recent studies noting a series of key architectural differences compared to humans^12–14^.

Indeed, recent 3D imaging and topological mapping of the murine LN^15,16^ have described a mesh-like network composed of fibroblastic reticular cells (FRCs) that provide the homing zone for T cells and may mediate B cell homeostasis^17^. Moreover, Huang and colleagues recently demonstrated the murine LN capsule to be encased by a network of sensory neurons capable of neuroimmune transcriptional regulation^18^. The delivery of antigens^19^, inflammatory mediators, and even larger molecules^20^ to the LN T cell zone involves the transport of lymph initially filtered through the subcapsular sinus, followed by passage along an elaborate lymphatic conduit system^21^. Similarly, Ogembo et al.^13^ and Buckley et al.^22^ have elegantly described SIRPα, FHOD1, HLA-DR, CD36 (platelet glycoprotein IV), CD71 (transferrin receptor) and CD8α as cell markers expressed by littoral cells (LCs) lining the splenic sinusoids. LCs exist exclusively in human SPL and are not observed in rodents or even non-human primates^13,22^. However, human tissue studies have historically been limited to two-dimensional (2D) tissue cross-sections stained for low-parameter combinations of markers, highlighting the need to characterize the human SPL and LN using a combination of highly multiplexed 2D and three-dimensional (3D) modalities.

### Multiplexed 2D-imaging of immune, lymphatic and blood vessel markers in human SPL and LN

To extensively profile the cellular architecture of immune cells along with lymphatic and blood endothelial cells (LECs and BECs, respectively) within the human SPL and LN, antibodies were first validated by immunohistochemistry (IHC), demonstrating the expected organ-specific staining patterns within formalin fixed paraffin embedded (FFPE) tissue sections (Extended Data Fig. 1). SPL and LN tissue sections were then stained with a 29-marker panel (Supplemental Table 1) for highly multiplexed imaging using CODEX (CO-Detection by indexing)^23,24^. Visualizing the expression patterns (Extended Data Fig. 2) for immune cell type/subset-defining markers^25,26^ [B cell (CD20, CD21), follicular dendritic cell (FDC; CD21, CD35), T cell (CD3ε, CD4, CD8α, CD45RO), neutrophil (CD15), and macrophage (CD68, CD163)] delineates the organ-specific structures comprised of repeating functional units (i.e., lymphoid follicles). When combined with the endothelial markers [LYVE1, podoplanin (PDPN), and PROX1 for LECs^25,27^; CD31 and CD34 for BECs^28,29^], one is able to appreciate the intricate cellular architecture of “normal”/non-diseased LN and SPL (Extended Data Fig. 3-5).

Contrary to observations from single cell transcriptomic profiling of murine LN^4,5^, our results demonstrate that LYVE1 and PDPN expression do not overlap in the human LN or SPL (Fig. 1). To substantiate these observations, a pair-wise pixel-based Pearson’s correlation analysis was utilized to generate averaged R values, providing a measure of signal correlation in LN and SPL (Extended Data Fig. 5e-f). Indeed, the LYVE1/PDPN marker pair had little to no overlap (LN, R = 0.028; SPL R =0.0), in contrast with marker pairs such as CD4/CD3 (LN, R = 0.69; SPL, R = 0.48) and CD21/CD35 (LN, R = 0.57; SPL, R = 0.61) which were moderately correlated, as expected^30^.

**Figure 1.**
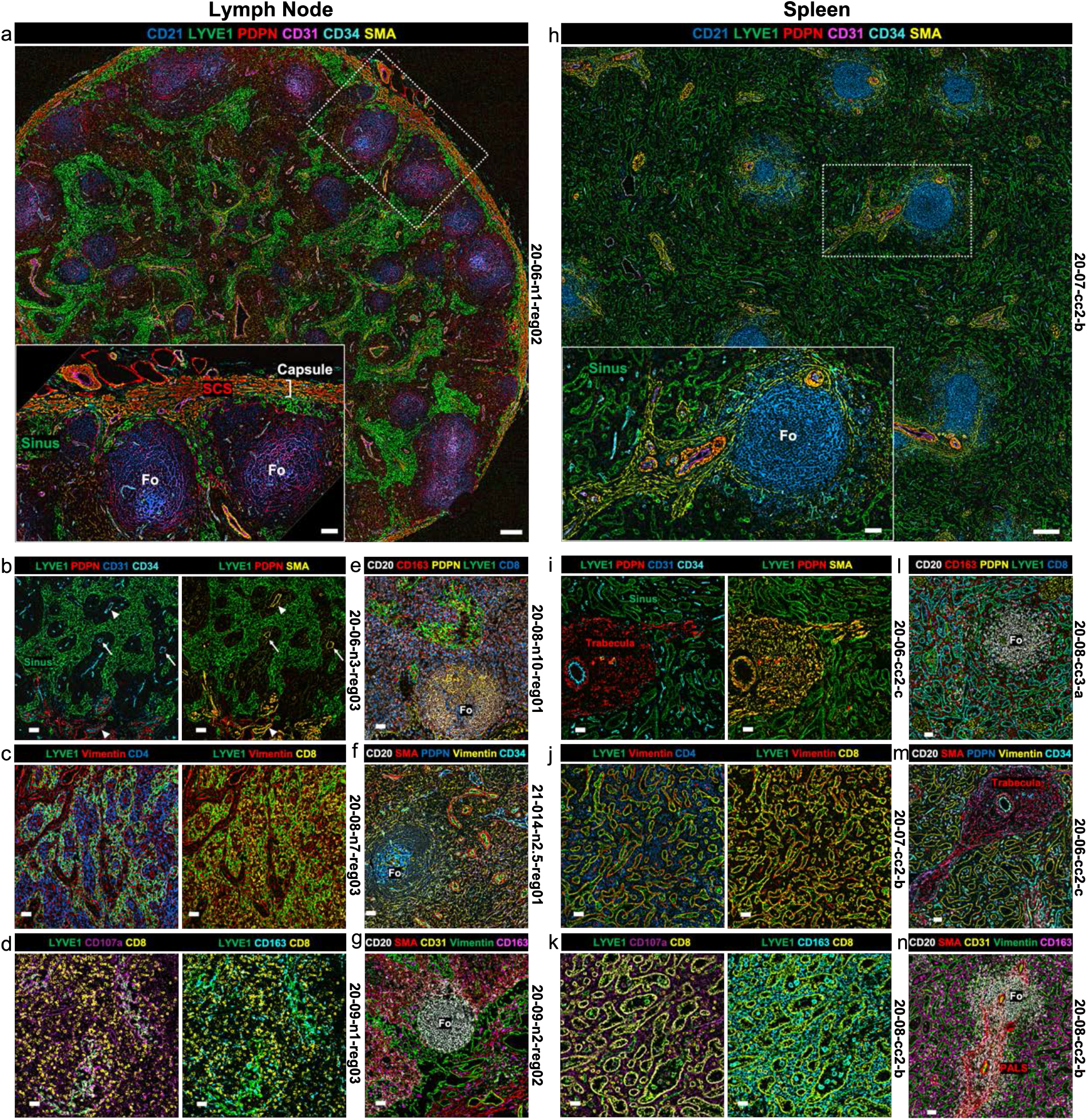
Immune, LEC, BEC, and stromal cell organization in human LN and SPL. Representative CODEX images of **a-g**, LN and **i-n**, SPL. **a**, LN cross-section (scale bar 690μm) with annotated inset (scale bar 180μm) showing capsule, subcapsular sinus (SCS), follicles (Fo; CD21+PDPN+), and lymphatic endothelium within the sinus (LYVE1+). The LN showed normal microarchitecture with respect to sinus distribution and follicular meshwork formation. **b**, CD31+CD34+SMA+ high endothelial venule (HEV, white arrow) within the LYVE1+ sinus along with PDPN+ veins (white arrowheads). **c**, CD4+ (left) and CD8a+(right) interfollicular T cell zone with Vimentin+ stroma. **d**, CD107a cytotoxic lymphocytes and CD163 macrophages within the interfollicular space. **e**, CD20+ B cells enrich the PDPN+ follicular dendritic cell (FDC) meshwork within a Fo with CD8a identifying the T cells within the interfollicular space along with CD163+ macrophages. **f**, CD20+ B cells enrich the PDPN+ Fo with SMA highlighting proximal arteries and CD34 highlighting venules. Vimentin illustrates background stromal architecture. **g**, CD20 identifies a Fo with CD163 highlighting interfollicular macrophages and with SMA, Vimentin, and CD31 highlighting stroma and endothelium. **h**, SPL cross-section (scale bar 690μm) with annotated inset (scale bar 180μm) showing CD21+PDPN-Fo, LYVE1 + venous sinusoids (Sinus), with CD31+CD34+SMA+PDPN+ highlighting the arterioles and periarteriolar lymphoid sheaths (PALS). **i**, CD34+PDPN+SMA+ trabecula surrounded by LYVE1 + venous sinusoids. **j**, CD4+ T cells (left) within the cords of Billroth; CD8a+ LCs with LYVE1+ lymphatic endothelium and LYVE1-CD8a+ T cells within the cords of Billroth (right). **k**, Cords of Billroth with CD107a+, CD8a+/− cytotoxic lymphocytes and CD163+ macrophages arranged between the CD8a+ LCs and LYVE1+ sinusoidal lymphatic endothelium. **l**, CD20+PDPN-Fo surrounded by CD8a+ LC and LYVE1+ sinusoidal lymphatic endothelium with CD163+ macrophages in the cords of Billroth. **m**, SMA+CD34+ trabecula with Vimentin+ highlighting background stroma and only rare CD20+ B cells and absence of PDPN+ FDCs. **n**, White pulp with tangentially sectioned CD20+ Fo and adjacent SMA+ PALS. Vimentin highlights stroma and CD163+ macrophages in cords of Billroth. **b-g and i-n**, Scale bars, 180μm. Donor ID, tissue, and region defined to the right of each panel.

PDPN and smooth muscle actin (SMA) appear to define the LN subcapsular sinus^31^ (SCS), as depicted in Fig. 1a with annotations (inset) of the LN including lymphoid follicles and lymphatic endothelium^6,31^. Indeed, our results illustrate the expected expression patterns for LYVE1^32,33^, which identifies the complex network of LN LECs^15,16^ (Fig.1a,b,c,d,e). Consistent with previous studies^34,35^, PDPN+ cells are observed within LN follicles (Fig. 1a,e,f; PDPN/CD21 R = 0.21, PDPN/CD35 R = 0.24, Extended Data Fig. 5e). Within SPL, we are able to visualize follicles, periarteriolar lymphoid sheath (PALS), trabeculae, and associated trabecular arteries positioned amongst the intricate network of venous sinuses, as shown in Fig. 1h-i and annotated within the inset. Interestingly, LYVE1 labels CD8α+ LCs lining the venous sinusoids^13,36^ (Fig. 1j,k). In contrast with LN, PDPN expression is absent in SPL follicles (Fig 1h,l; PDPN/CD21 R = 0.02, PDPN/CD35 R = 0.08, Extended Data Fig. 5f). Finally, PDPN is observed surrounding large arteries in both organs (Fig. 1a,h), and specifically in splenic trabeculae (Fig. 1i,m).

Vimentin is observed throughout the LN sinuses and endothelium labeled by CD34, SMA, and CD31 (Fig. 1c,f,g). In the SPL, vimentin partially overlaps with LYVE1 (R = 0.44), most notably in splenic sinusoids identifying CD8α+LCs^13^ (Fig. 1j). The CD8αβ heterodimer is a canonical T cell marker^37^ and as expected, CD8α labels individual T cells scattered across the LN (Fig. 1c,d,e and Extended Data Fig. 2b). However, within the SPL, CD8α also defines the LCs of the splenic sinuses, in line with previous reports^13^, overlapping with vimentin and LYVE1 (Fig. 1j,k,l). CD107a (lysosome-associated membrane glycoprotein 1 [LAMP1]), which marks degranulation on Natural Killer (NK) cells and CD8^+^ T cells^38^ along with leukocyte adhesion to vascular endothelium^39^, and the monocyte/M2 macrophage marker CD163 are observed throughout the LN sinus network (Fig. 1d,g) while in SPL, both markers localized to the cords of Billroth between venous sinuses within the red pulp (Fig. 1k,n). To corroborate the representative images in Figure 1, we provide additional CODEX datasets both from independent donors and within multiple common coordinate framework (CCF)^40,41^-defined tissue regions for both LN and SPL (Extended Data Fig. 3-5), and complete CODEX datasets are publicly available on the HuBMAP data portal (Extended Data Table 1).

### Cell clustering and quantitation in 2D

For a subset of donors, we utilized a 16 marker spatial X-shift clustering^42^ to segregate and classify various cell populations in both LN and SPL (Fig. 2a). We then annotated cell clusters based on canonical marker expression and tissue localization. As anticipated^43^, T cells account for approximately one-half of all cells in LN; this, in contrast with approximately 10% of all cells in SPL. The majority of T cells in either tissue are CD8^+^ (Fig. 2b), as expected^44–46^. The CD15^+^neutrophil population provides a further distinction between the two organs comprising approximately 1%of cells in LN versus 30% in SPL, which is in line with the literature^47–49^. A color coordinated Voronoi tessellation image provides a visual illustration of annotated cell populations organized within and around a follicle in each tissue (Fig. 2c). Specifically, B cells and FDCs localized within the follicle, surrounded by marginal zone B cells, which were more pronounced in LN than in SPL, in agreement with prior reports^50^. Follicles were further surrounded by a distributed mixture of the other cell clusters.

**Figure 2.**
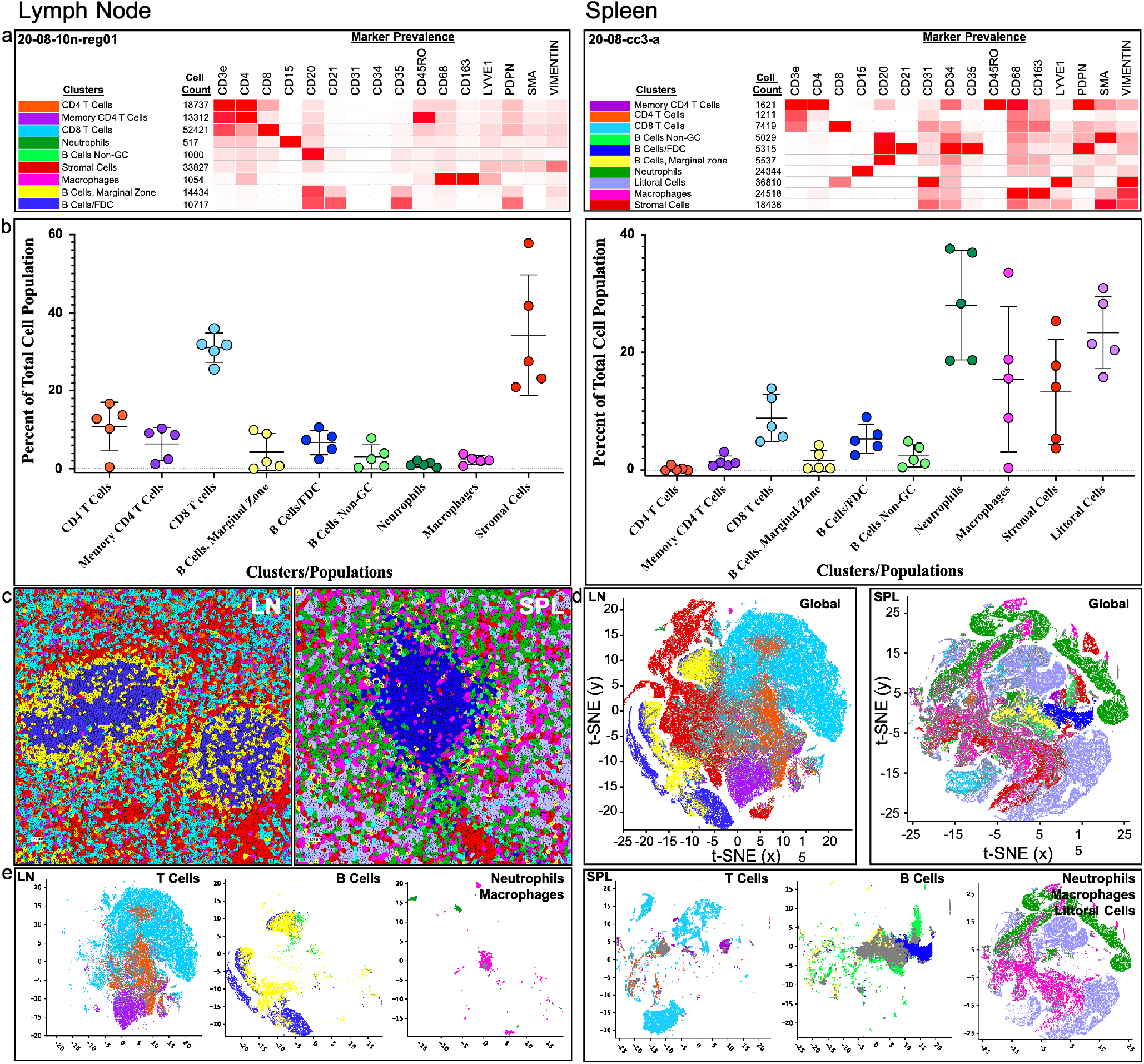
Cell clustering, segmentation, and t-SNE visualization. Analysis of CODEX data from LN (*left*, n=5) and SPL (*right*, n=5) with identical color coding for each cluster throughout and Littoral cells included as an additional cluster for SPL. **a**, Supervised X-shift clustering with heatmap illustrating protein markers that define each cluster and cell counts for each cluster with donor IDs in upper left corner. **b**, Cell counts for each supervised cluster are plotted as the percent of the total cell population for each donor image. **c**, Clustered Voronoi plot of follicles prominently highlighting the three supervised B cell populations in the LN and SPL. **d**, Global high dimensionality reduction of clustered data set using t-SNE (t-Distributed Stochastic Neighbor Embedding). **e** Three gated t-SNE plots for T (upper), B (middle), as well as neutrophil, macrophage, and/or Littoral cell (lower) populations. Analyses performed with Akoya CODEXMav plugin to NIH Fiji using default algorithm parameters.

We next performed a nonlinear high dimensionality reduction using t-Distributed Stochastic Neighbor Embedding (t-SNE)^51^ to project all clustered populations defined in Fig. 2a into low-dimensional space (Figs. 2d). Foremost, these renderings highlight the dramatic differences in T cell and neutrophil population frequencies within the two organs, which is further illustrated by the gated graphs displaying individual total T cell, B cell, and neutrophil/macrophage populations in the LN and SPL (Fig. 2e). Fig. 2e (far right) also demonstrates the proximity of the macrophage population in the cords of Billroth (Fig.1k,n) with the LCs lining the splenic sinuses. Cellular clustering, Voronoi images and global as well as gated t-SNE images for four additional LN and SPL are provided in Extended Data Figures 6 and 7, respectively.

### 3D-imaging of perivascular innervation in SPL and LN

To visualize vascular innervation of the human LN and SPL in 3D, we employed multiplexed, three-color light sheet fluorescence microscopy (LSFM)^52,53^ on tissues cleared using CLARITY-based methodology^54,55^. An example of a pre- and post-cleared intact human LN is shown in Extended Data Figure 8a. Extensive 3D perivascular innervation is observed on CD31+ endothelial cells within vessels of human LN (Fig. 3a-d) and SPL (Fig. 3e-h). CD31 specificity is visualized at the individual endothelial cell level in the SPL (Extended Data Fig. 8b,c). Staining with the pan-neuronal marker PGP9.5 reveals a complex matrix of neural processes surrounding CD31+ vasculature of the LN (Fig. 3b-d) and SPL (Fig. 3f-h) located within the *nervi vasorum* of the *tunica externa* layer^56^ of the vessel wall (Extended Data Fig. 8d). *Nervi vasorum* staining is also observed in SPL for the peripheral sympathetic neuronal marker tyrosine hydroxylase (TH) (Extended Data Fig. 8e, Supplemental Video 1). Interestingly, choline acetyltransferase (ChAT), an enzyme required for production of the neurotransmitter acetylcholine within parasympathetic neurons^57^, colocalizes with a subset of the PGP9.5+ perivascular axons in the LN (Fig. 3a-d) and SPL (Fig. 3e-h). This is in contrast with the previous literature^58^, including recent IHC data from human cadaveric specimens where PGP9.5^+^ staining was documented in the absence of ChAT^7^ as well as 3D imaging of cleared SPL from mice where ChAT^+^ nerve fibers were not detected^59^. However, our intensity-based volumetric segmentation demonstrates that 21% of the PGP9.5+ neuronal processes also contain ChAT in the LN evaluated, and similarly, 22% contain ChAT in SPL (Fig. 3c,g). Renderings of sequential optical slices through the Z dimension provide 3D visualization of perivascular innervation within human LN (Supplemental Video 2) and SPL (Supplemental Video 3), demonstrating that PGP9.5, ChAT, and TH primarily label neuronal processes on large vessels in both organs.

**Figure 3.**
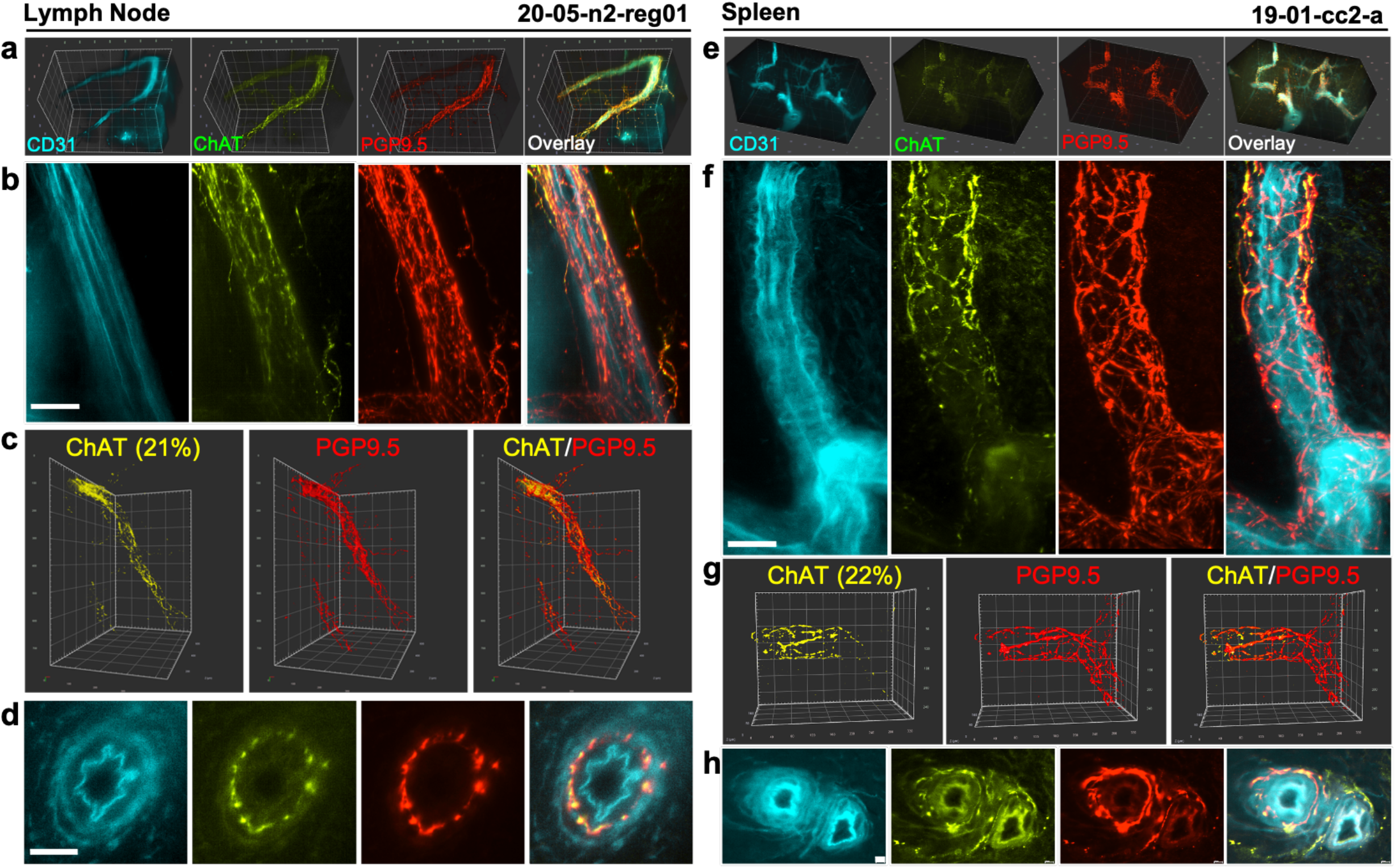
Perivascular innervation of human LN and SPL imaged with 3D LSFM. Multicolor LSFM imaging of CD31+ vessels with ChAT and PGP9.5 innervation in **a-d**, LN and **e-h**, SPL. **a, e**, 3D renderings of 0.5 mm^3^ (LN) and 1 mm^3^ (SPL) tissue volumes. **b, f**, Max intensity projection of vessels expressing CD31 surrounded by extensive PGP9.5-positive innervation with partial ChAT-positive neuronal processes in LN (100 μm) and SPL (50 μm). **c, g**, 3D renderings of perivascular neuronal network illustrating a subset of ChAT+ axons with the percent of total PGP9.5+ axons expressing ChAT determined using semi-automated, intensity-based volumetric segmentation with the Arivis software package. **d**, **h**, Cross-section through LN (50 μm) and SPL vessels (10 μm) demonstrating co-localization of ChAT and PGP9.5 neural components within the *tunica externa*. Donor ID, tissue, and region are defined at the top.

LSFM imaging of LN further identifies perivascular innervation of large vessels expressing the pan-neuronal marker β3-tubulin as well as an extensive mesh-like vessel network expressing both β3-tubulin and neuronal growth-associated protein (GAP43, or neuromodulin)^15,16,60,61^ (Fig. 4a-c; Supplemental Video 4). Indeed, in comparison to 2D analysis, LSFM affords the ability to better appreciate these complex structures in 3D. Indeed, in Extended Data Fig. 9a, we provide a close examination of high-resolution 3D max-projection images. We also observe β3-tubulin+ nerve bundles adjacent to blood vessels (Fig. 4b; Supplemental Video 5), along with a single GAP43^+^ axon within a vessel labeled by β3-tubulin (Extended Data Fig. 9b).

**Figure 4.**
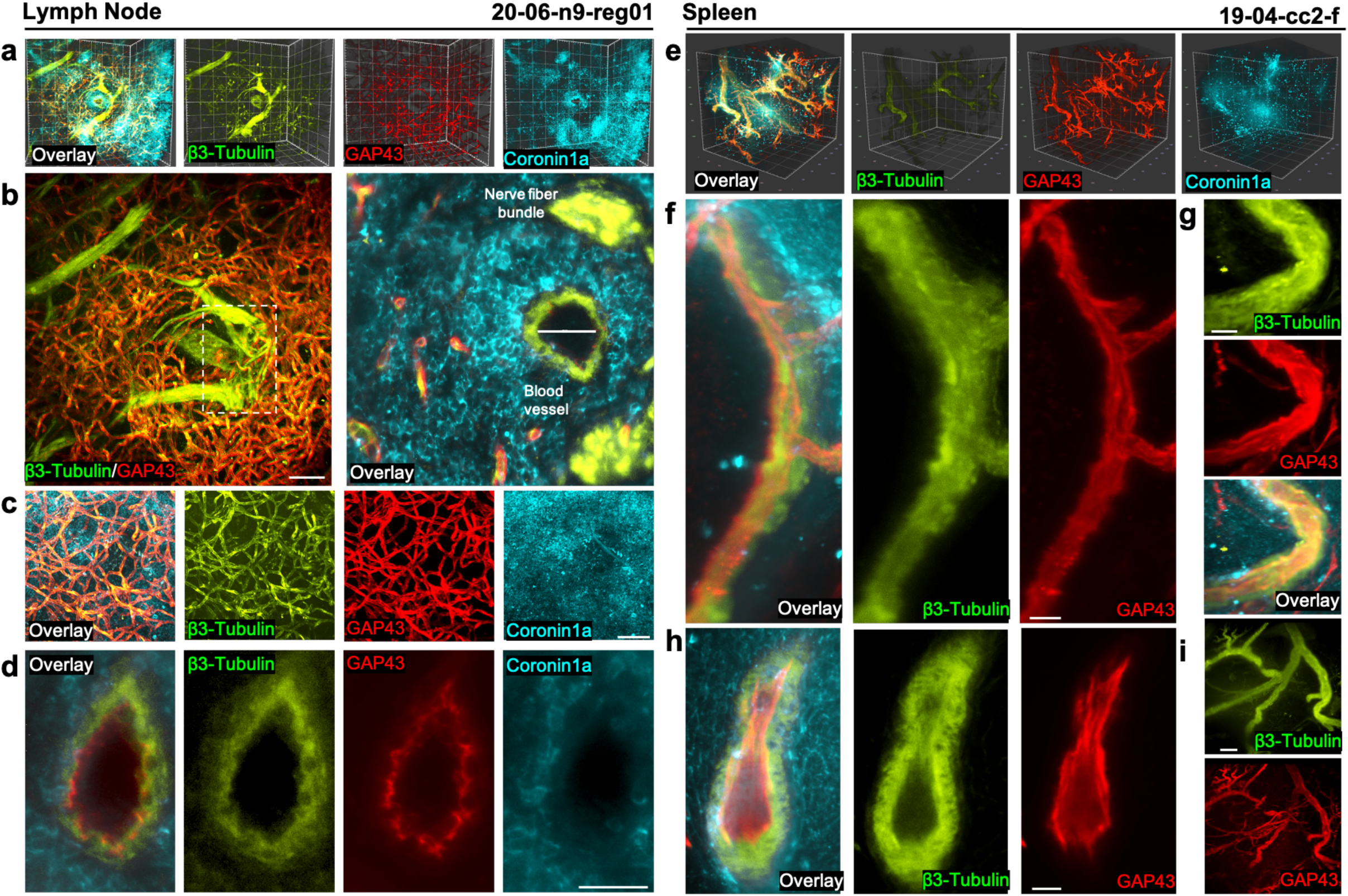
Neural markers along vessels of human LN and SPL. Multicolor LSFM imaging of β3-tubulin and GAP43 innervation with Coronin-1A+ leukocytes in **a-d**, LN and **e-i**, SPL. **a**,3D rendering of 0.5 mm^3^ LN tissue. **b**, Max intensity projection (*left)* of LN large vessel and nerve bundles expressing β3-tubulin woven between an extensive β3-tubulin+GAP43+ small vessel network (100 μm) with inset showing zoomed in cross-section (*right)* with blood vessel diameter annotated (60 μm). **c**, Wide-field max intensity projection showing Coronin-1A+ leukocytes among a complex network of β3-tubulin+GAP43+ small diameter vessels (100 μm). **d**, LN vessel cross-section demonstrating localization of β3-tubulin to the *tunica externa* and GAP43 to the *tunica interna* of the vascular wall (50 μm). **e**, 3D rendering of 1 mm^3^ SPL tissue. **f, g**, Max intensity projection of SPL vessels expressing β3-tubulin within the wall and GAP43 within the internal lining (20 μm). **h**, Max intensity projection showing β3-tubulin+GAP43+ large and GAP43+ small vessels (100 μm). **i**, SPL vessel cross-section demonstrating localization of β3-tubulin to the *tunica externa* and GAP43 to the *tunica interna* of the vascular wall (20 μm). Donor ID, tissue, and region are defined at the top.

As expected, the leukocyte marker Coronin-1A^62^ is detected throughout the LN (Fig. 4a-c; Supplemental Video 5). In contrast, Coronin-1A expression is less homogenous in SPL, observed both on single cells and on cells clustered around large vascular structures identified by perivascular nerves expressing β3-tubulin (Fig. 4e). Within the SPL, β3-tubulin identifies large vascular processes, whereas GAP43 identifies both large and small vessels (Fig. 4e-i), corresponding either to BEC or efferent LECs^63^. Interestingly, immunolabeling with GAP43 demonstrated staining localized to the internal elastic membrane or smooth muscle layer of large blood vessels in both LN (Fig. 4d) and SPL (Fig. 4h). These GAP43 data clearly define filamentous neuronal processes existing on the internal aspect of large vessels (Extended Data Fig. 8d), unexpected from conventional *nervi vasorum* location of vascular innervation^7^. To our knowledge, this localization of GAP43 expressing cells has not previously been reported in human LN or SPL vessels.

### Differential expression of perivascular GAP43 and β3-tubulin in LN and SPL

A detailed review of our 3D LSFM data reveals a novel observation involving differential expression of β3-tubulin and GAP43 along discrete segments of large vessels in both LN and SPL (Fig. 5a-g). Specifically, within LN, select segments of large vessels display a reduction or absence of protein expression for either GAP43 (Fig. 5a) or β3-tubulin (Fig. 5b,c) as quantified using an intensity-based volumetric segmentation (Extended Data Fig. 10a,b).

**Figure 5.**
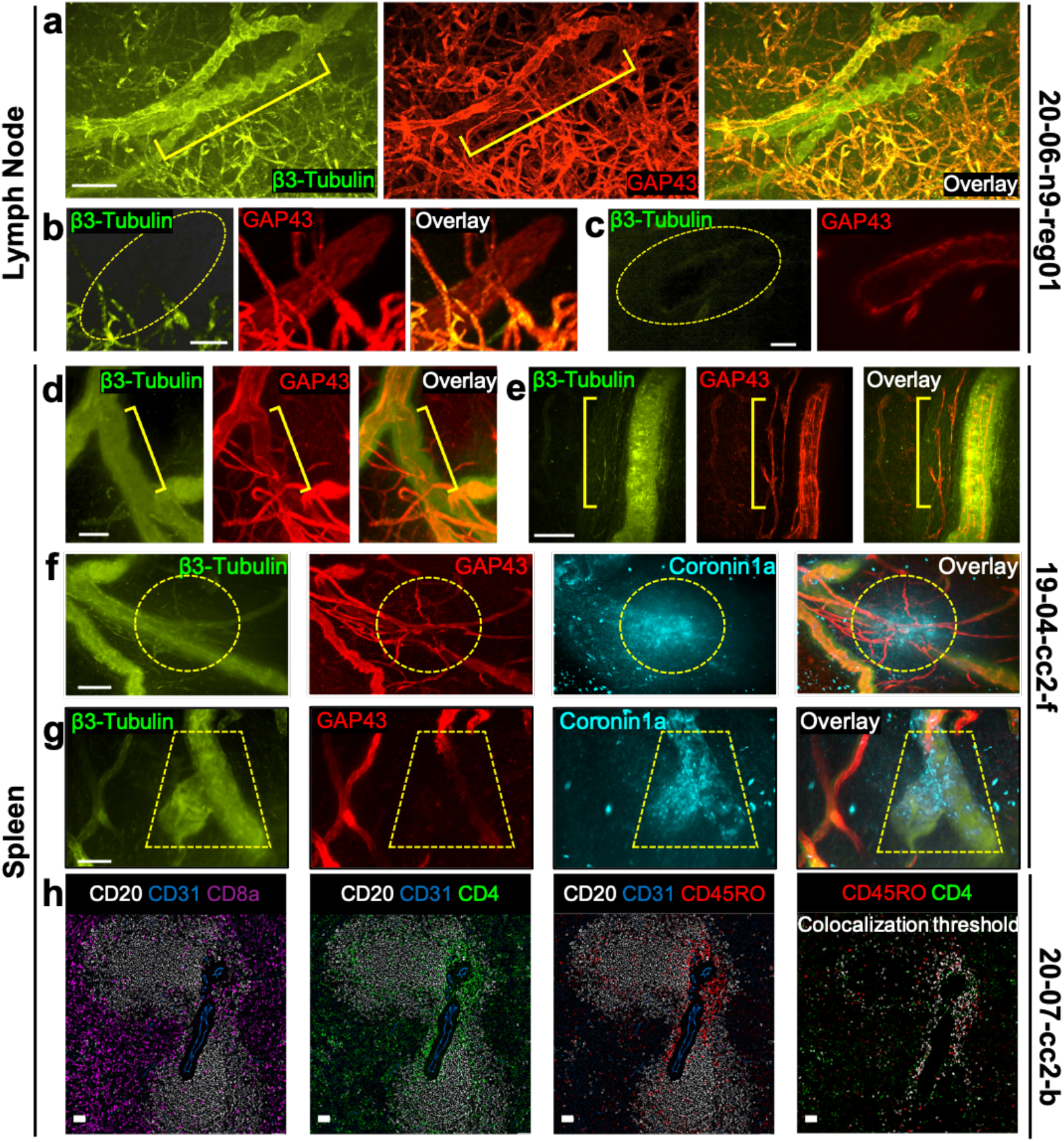
Differential expression of β3-tubulin and GAP43 along discrete segments of large blood vessels in human LN and SPL. Multicolor LSFM imaging of β3-tubulin and GAP43 innervation in LN (**a-c**) and SPL with Coronin-1A+ leukocytes (**d-g**). **a**, Max intensity projection showing large β3-tubulin+ vascular segment with a region of decreased GAP43 expression (yellow bracket; 100 μm). **b**, 3D rendering (40 μm) and **c**, longitudinal cross-section of LN vessel (20 μm) demonstrating absence of β3-tubulin (dashed circle) where GAP43 expression is present. **d, e**, Max projections (50 μm) show a β3-tubulin+GAP43-vessel (**d**) and a β3-tubulin-GAP43+ vessel (**e**) indicated with yellow brackets in SPL. **f, g**, Max intensity projection of SPL vessels showing regions of continuous β3-tubulin, reduced GAP43, and clustered Coronin-1A expression (dashed outlines; 50 μm). **h**, Multiplexed CODEX image of two CD20+ splenic follicles with a CD31+ central artery passing through a PALS region demonstrating a clustering of CD4+CD45RO+ memory T cells around the artery, in contrast to a perifollicular distribution of CD8a+ T cells (180 μm). Donor ID, tissue, and region are defined to the right.

Similarly, GAP43 protein expression is notably diminished along segments of large vessels within the SPL, in the context of contiguous β3-tubulin expression (Fig. 5d,e). Figure 5d also reaffirms the observation that within the SPL, GAP43 detects neuronal elements on both large and small vessels, while β3-tubulin is primarily found on cells along large vessels (Fig. 4e,i). Notably, diminished expression of GAP43 along segments of large vessels in the SPL appears to associate with densely clustered populations of Coronin-1A+ cells (Fig. 5f,g, Supplemental Video 6), as supported by quantitative volumetric segmentation (Extended Data Fig. 10c,d). To further this observation, we utilized CODEX to examine a region of the SPL that included a PALS with CD45RO+ T cells surrounding blood vessels (Fig. 5h and Extended Data Fig. 11) and noted that these T cell clusters may potentially correspond to the Coronin-1A+ cells observed in LSFM.

## Discussion

Herein we provide novel 2D and 3D characterization of the immune, endothelial, and neuronal cell networks within the normal human LN and SPL. Among key observations, LYVE1 and PDPN do not overlap in either tissue, as supported by pairwise pixel-based quantitation. Specifically, PDPN and SMA label the LN subcapsular sinus with LYVE1 also staining the lymphatic endothelium within the LN interior. Importantly, studies using a PDPN+ cell sorting strategy to isolate LEC from murine LN prior to single cell transcriptomics, may fail to observe the LYVE1+ population of cells^4,5^. Building on a seminal studies of LCs^13^ and endothelial cells^27^, we report that LYVE1 labels LCs lining the SPL sinusoids, along with vimentin and CD8a, while PDPN is expressed in central arteries and trabeculae. LCs do not exist in rodents or non-human primates, emphasizing the importance of performing these cell mapping studies in human tissue^13^. The abundance of macrophages expressing CD163 (hemoglobin scavenger receptor) within the splenic cords of Billroth is in line with the existing literature and supports this organ’s well-established function as the primary site for filtration of aged or damaged red blood cells from circulation^13,64^. Neighborhood analysis using our CODEX data, coupled with population cell counts, delineates the disparate composition of T cell and CD15+ neutrophil populations in the LN versus the SPL, in line with prior reports^44–49^. In particular, we observe a much higher proportion of CD3+CD8a+ T cells in the LN compared with SPL. In addition, we note a smaller population of CD15+ neutrophils in the LN versus the SPL, corroborating the existing literature^65,66^.

Our LSFM data highlight the power and utility of 3D imaging to observe intact tissue volumes, consisting of thousands of optical sections from any vantage point, providing new insights into human tissue organization that may be masked by max projection, single-section 2D views, or 3D stitching of 2D images. This effort, which applied three-color immunofluorescent staining to human secondary lymphoid organs, represents an advance on prior LSFM imaging studies^52,53,67–69^. Moreover, given HuBMAP’s use of a CCF^40,41^, the images presented herein can be mapped back to their physical location within the intact organ. Within both LN and SPL, the PGP9.5, β3-tubulin, ChAT, and TH markers identify an extensive network of perivascular neuronal processes localized to the *nervi vasorum* of the *tunica externa*, while GAP43 appears to localize to the internal elastic membrane or smooth muscle layers of the artery^70^. We further observe that nerves expressing GAP43 and β3-tubulin associate with an extensive network of small vessels within the LN. Moving forward, there is a need to corroborate our interpretation that LN LYVE1 staining, visualized in 2D via CODEX, may correspond with this vessel network.

The leukocyte marker Coronin-1A identifies scattered single cells in LN imaged by LSFM. In contrast, Coronin-1A+ cells are both scattered and clustered around vascular structures in SPL. Interestingly, accumulation of leukocytes around SPL blood vessels (PALS) appears associated with reduced expression of GAP43. In support of this observation, 2D SPL CODEX images reveal similar clustering of CD4+ T cells in PALS. Future work will require functional assays to determine if regions lacking GAP43 correspond to sites for neurovascular regulation of immune cell extravasation and navigation^71,72^.

The data presented herein showcase the unique staining patterns of immune, endothelial, and neural markers within human secondary lymphoid organs. Given that most human immune responses occur in secondary lymphatic organs and not peripheral blood^30^, there is a clear need for additional studies using human tissues to gain insights on how neuroimmune interactions modulate cellular activity across the lymphatic system. Importantly, the current 3D cell mapping effort, together with complementary datasets and tools generated across the HuBMAP Consortium^41,73–77^, among others (e.g., Human Cell Atlas^78^), also set the stage for future studies to determine how these pathways may be dysregulated in various disease settings, including autoimmunity, sepsis, cancer, and viral infection. We envision that novel therapies targeting the neurovascular sites identified herein could potentially be developed to specifically modulate immune function within the human SPL and LN.

## Methods

### Donor Acceptance Criteria

SPL and LN were acquired from 14 non-diseased organ donors (Extended Data Table 1) through the Human BioMolecular Atlas Program (HuBMAP)^3^ Lymphatic System Tissue Mapping Center (TMC) at the University of Florida, via a nationwide organ procurement network. Prior to organ recovery, written informed consent for research is provided by the donor families. Criteria for donor acceptance by the HuBMAP Lymphatic System TMC at the University of Florida are available on the HuBMAP site (https://www.protocols.io/view/donor-acceptance-criteria-for-tmc-florida-zurich-h-bipykdpw) and as follows: donors less than 7 days on a ventilator, tissue received within 24 hours of recovery, and donor body mass index (BMI) between the 5^th^ and 95^th^ percentile as defined by the United States Centers for Disease Control and Prevention (CDC). Contraindications for acceptance include: more than four blood transfusions, autoimmune disease, chromosomal abnormalities, positive culture for bacterial or viral meningitis, or methicillin-resistant *Staphylococcus aureus* (MRSA), sepsis, observable infarction, rupture, SLP or LN deflation or gross abnormality, immune deficiency disease, cancer, infection including severe acute respiratory syndrome-coronavirus-2 (SARS-CoV-2), and splenomegaly.

### SPL and LN Case Processing and Quality Control (QC)

Human SPL and mesenteric LN were processed within 16 hours of cross clamp. Residual fat or connective tissues were removed from each LN and size (mm) recorded. LN > 1 cm^2^ were bisected to 1 cm^2^. LN < 1 cm^2^ were left intact. Intact spleens were photo documented with hilum to left, inferior end at top. Cross-sectional slabs were cut 1 cm thick, and a 1 cm wide strip was cut through the center of each slab with the superior pole surface facing up and the hilum serving as the point of origin. 1 cm^3^ blocks were dissected from each strip with all cuts photo documented for location and orientation. Tissue cassettes were placed in at least 20 volumes of 4% PFA (i.e., 20 mL 4% PFA per 1 mL tissue) for 20-24 hours, transferred to 70% ethanol, and paraffin embedded in an automated tissue processor (Sakura VIP) within 3 days. SPL and LN were sectioned to 5 μm and stained with hematoxylin and eosin (H&E) and evaluated by an independent pathologist to assess organ normality.

### Immunohistochemistry

IHC was performed according to HuBMAP Lymphatic System TMC standard operating procedures (SOPs) as detailed (https://www.protocols.io/view/dab-staining-of-ffpe-slides-bdiei4be). Formalin fixed, paraffin embedded (FFPE) human SPL and LN were sectioned to 5 μm, mounted on SuperFrost Plus (Fisher Scientific) microscope slides and air dried overnight. Slides were deparaffinized twice in xylene and 100% ethanol for 5 minutes, peroxidase blocked with 3% peroxide in methanol for 10 minutes followed by serial ethanol washes (95%, 75% and 50%) each for 5 minutes. Finally, slides were washed in deionized water twice for 5 minutes. Antigen retrieval involved microwaving at 50% power for 7 minutes in 10 mM sodium citrate buffer pH 6.0 followed by an 18 minutes incubation in the hot citrate. Slides were then rehydrated with 50mM Tris buffered saline pH 7.6 with 0.1% Tween (TBST) for 5 minutes in a humidified chamber. Diaminobenzidine (DAB) staining was performed using VECTASTAIN Elite ABC Kit (peroxidase-HRP) kit (Vector Labs) and IMPACT DAB (peroxidase substrate). Avidin and biotin were sequentially blocked using Avidin/Biotin Blocking Kit (Vector Labs) for 20 minutes, followed by TBST rinse and incubation in 2% normal serum block (in TBST) for 1 hour. Primary antibodies were diluted in Antibody Diluent Reagent Solution (Life Technologies; Supplemental Table 1), applied to the slide, covered with a plastic coverslip and incubated overnight at 4°C in humidified chamber. Slides were incubated with diluted biotinylated secondary antibodies (5 μl antibody, 15 μl serum and 1 mL TBST) for 30 minutes at room temperature followed by addition of ABC Reagent for 30 minutes. Diluted DAB reagent was then added, development observed microscopically and reaction stopped in tap water. Slides were then counterstained with hematoxylin followed by alcohol and xylene dehydration and cover-slipped.

### CODEX Antibody Conjugation and Validation

Commercially available oligonucleotide conjugated antibodies were purchased from Akoya Biosciences and used at a standard dilution of 1/200 (Supplemental Table 1). Akoya CODEX antibody validation consists of comparing its oligo-conjugated antibodies with dye-conjugated antibodies to assure equivalent staining pattern, antibody titration for optimal signal, and single and multiplex staining for consistent staining patterns (https://www.akoyabio.com/wp-content/uploads/2022/01/Phenocycler_Technical-Note_Validation-of-Commercial_DN-00140.pdf). For custom conjugation, purified, carrier-free, antibodies are purchased from reputable vendors, providing validation data including IHC-Paraffin and western blots. All antibodies, including commercially available CODEX antibodies are validated by IHC in our lab using the tissue of interest. Barcodes are assigned to antibodies based on the abundance and intensity of the protein expression. Less abundant antigens are assigned to fluorescence channels having lower natural autofluorescence, namely Cy5 and AF750. Conversely, antibodies for highly expressed antigens can be assigned to ATTO550 where tissue autofluorescence is higher. Conjugations were performed with commercially available CODEX Conjugation Kits (Akoya Biosciences) according the manufacturer’s recommendation with the following modifications. Briefly, for each antibody, a 50 kDa molecular weight cut-off (MWCO) filter (Amicon Ultra) was washed with Filter Blocking Solution, centrifuged at 12,000 xg for 2 minutes and emptied of all liquid. To concentrate the antibody, 50 μg of protein, as determined by Nanodrop absorbance, was diluted in 100 μl PBS, added to the filter and centrifuged at 12,000 g for 8 minutes. To initiate the disulfide reduction reaction, Antibody Reduction Master Mix was added to the filter, vortexed for 2-3 seconds, incubated at room temperature for no more than 25 minutes followed by buffer exchange. A unique barcode was prepared for each antibody (Supplemental Table 1) by reconstituting in 10 μl distilled water, adding 210 μl of the Conjugation Solution with incubation at room temperature for 2 hours. 5-10 μl of antibody solution was reserved for protein electrophoresis, and the filter washed three times with Purification Solution. Newly conjugated antibodies were collected by centrifugation. Custom conjugated antibodies were titrated (1/200, 1/100, 1/50) and validated by comparing the staining patterns of each conjugated antibody alone in CODEX against IHC DAB staining before being included in multiplex experiments. Newly conjugated antibodies were also stained alongside positive and negative controls in the CODEX platform as described (https://www.protocols.io/view/hubmap-tmc-uf-validation-of-custom-conjugated-anti-bkpzkvp6).

All antibodies, including commercially available CODEX antibodies are validated by IHC in our lab using the tissue of interest. Barcodes are assigned to antibodies based on the abundance and intensity of the protein expression. Less abundant antigens are assigned to fluorescence channels having lower natural autofluorescence, namely Cy5 and AF750. Conversely, antibodies for highly expressed antigens can be assigned to ATTO550 where tissue autofluorescence is higher.

### CODEX Staining

Oligonucleotide barcoded antibody staining of tissue sections mounted on cover slips (n=1 section per tissue region per donor) was performed using a commercially available CODEX Staining Kit according to the manufacturer’s instructions for FFPE tissue (Akoya Biosciences) and as recorded for HuBMAP Lymphatic System TMC SOPs (https://www.protocols.io/view/codex-antibody-staining-protocol-for-ffpe-tissues-bbsdina6). In brief, sample coverslips were heated to 55°C, cooled, deparaffinized and rehydrated. Antigen retrieval was performed using 1X Citrate pH 6.0 in a pressure cooker. Tissue coverslips were washed and equilibrated in CODEX Staining Buffer. Samples were incubated with barcoded antibodies (diluted as described in Supplemental Table 1) in CODEX blocking buffer for 3 hours in a humidity chamber at room temperature. After PBS washes, tissue sections were sequentially fixed with 1.6% paraformaldehyde (PFA), methanol (on ice) and Codex Fixative Reagent. Stained sample coverslips were stored for up to 2 weeks in CODEX Storage Buffer at 4°C. Barcoded fluorescent reporters corresponding to the barcoded primary antibodies were added to 96-well plates containing Codex Reporter Stock solution and nuclear stain, in groups of three (one of each wavelength) for each multiplex cycle (https://www.protocols.io/view/codex-preparation-of-reporter-96-well-plates-bc2riyd6).

### CODEX Image Acquisition

CODEX data were collected using CODEX Instrument Manager (CIM) software version 1.29, which automates image acquisition. During cycling, the CODEX instrument automatically adds the reporter solution to the coverslip, acquires fluorescent images, removes the reporters (DMSO/1X CODEX Buffer wash), and prepares the tissue for the next cycle. Images were acquired at 20X magnification using a Keyence BZ-X810 microscope with a metal halide light source, a Plan Apo λNA 0.75 20X air objective (Nikon), and the following emission filters (Chroma) and acquisition times: DAPI (358 nm, 10ms), TRITC (550 nm), CY5 (647 nm) and CY7 (750 nm) Alexa Fluor 750 (500ms), Atto 550 (350ms), and Alexa Fluor 647 (500ms). Keyence and image settings were multi-color z-stack, excitation light 100%, low photo bleach, Shadow 0, Highlight 255, gamma 1.0. The camera was set to mono and high resolution. Images were acquired in 13 cycles with the appropriate number of z-planes for the tissue of interest at 1.5 pitch: spleen 17 and lymph node 11. To address the consequences of and changes in autofluorescence during a CODEX experiment, a blank sample was imaged for each cycle providing a direct assessment of all background signal (autofluorescence). Images were 7×9 (3.77 × 3.58 mm). Raw images were saved in 14bit TIFF (*.tif) format.

### CODEX Image Processing

CODEX datasets were processed using the Akoya CODEX Processor 1.7.0.6. Multi-color z stacks acquired during each cycle were aligned by 3D drift compensation, and the best focal plane from each z stack was identified and used for quantitation. Specifically, there were four fluorescent channels with channel 1 (DAPI) used as the reference in cycle 2. Tiling was snaked by rows with 30% X and Y overlap, and 25 deconvolution iterations using the vectorial model. The segmentation parameters included: Anisotropic Region Growth, nuclear segmentation (DAPI), with a radius of 6, concentric circles 0, minimum/maximum cutoffs 0.02/0.99, relative cutoff 0.01, size cutoff factor 0.01 and inner ring size of 1.0. Advanced parameters included: tile processing (cycle alignment, background subtraction, deconvolution, extended depth of field), region processing (shading correction, tile registration, overlap cropping), stitching, full stitching, watershed segmentation, t-SNE report generation. Fluidic, temperature, and time dependent effects could occur between imaging cycles and were addressed with a using 3D phase correlation function. The calculated offsets were used in the alignment of all cycles as determined from the comparison of the reference cycle to the nuclear DAPI stain from the second cycle, for each cycle. Tile registration, from inaccuracies in camera positioning, were addressed using a Microscopy Image Stitching Tool (MIST) library^79^ that finds offsets by utilizing CUDA FFT phase correlation. Given the potential for variation of background signal across cycles, the normalized data were derived by subtracting background from each channel using both the first and last blank cycles. To address potential effects of exposure time, the Akoya software provides a ratio of signal exposure time to blank exposure time. For visualization purposes, some images had brightness subject to adjustment.

### Cell-Based and Neighborhood Analyses

We utilized the Akoya CODEX Multiplex Analysis Viewer (CODEX MAV; version 1.5.0.8 9) plugin in ImageJ (version1.53f51) for image analysis. The MAV software allows for image navigation, cell segmentation and biomarker quantification with clustering of cellular phenotypes accomplished by an X-shift clustering algorithm^42^ based on nearest neighbor density estimation. The cell types/clusters were manually annotated based on phenotype marker expression and displayed as an intuitive heatmap. Cell counts from each cluster from 5 LN and 5 SPL were then displayed as the percent of the total cell population. Voronoi analyses implemented as part of CODEX MAV were employed to illustrate separate B cell populations derived from marker-based annotation coupled with cellular localization. Neighborhood analyses for total data sets as well gated data subsets were displayed using dimensionality reduction and t-SNE visualization via the CODEX MAV software plugin.

Pixel-based pair-wise colocalization analyses were performed using the Coloc2 plugin (ImageJ/Fiji)^80^ to obtain Pearson’s correlation coefficients for respective protein marker pairings. Pearson correlation analysis, with R values near 1 indicating complete colocalization and R values near 0 indicating no colocalization of pixels above threshold intensity.

### Tissue Clearing

A CLARITY^81^ based protocol was used to clear tissues. All incubation and wash steps were performed with gentle rocking. Samples (< 1 cm^3^) were fixed in 40 mL of 4% PFA (Electron Microscopy Sciences) for 24 hours at 4°C, incubated for five days at 4°C in 40 mL of hydrogel monomer solution [4% acrylamide (Bio-Rad), 0.05% bis-acrylamide (Bio-Rad), 0.25% VA-044 Initiator (Wako Specialty Chemicals), 4% PFA, in PBS], then polymerized for 5 hours in 50 mL PBS at 37°C. Hydrogel embedded tissues were then cleared of lipids by incubating in 50 mL of clearing solution (Logos Bio) at 60°C, replacing clearing solution weekly until optically transparent. Cleared samples were washed in PBS with 0.1% Triton X-100 (PBST) for 48 hours at 37°C, replacing PBST wash buffer every 12 hours. The HuBMAP Lymphatic System TMC receives 1 cm^3^ pieces of SPL and LNs of varying sizes, ranging from 1 mm^3^ to 0.5 cm^3^. Samples are further portioned to test multiple immunolabeling combinations, generally we image 0.25 cm^3^ pieces of SPL, which is ~1% of the total spleen volume; whereas we image ~50% of each intact LN.

### Immunolabeling and LSFM 3D Imaging/Analysis

All incubation and wash steps were performed with gentle rocking. Cleared samples were immunolabeled by first incubating in 10 mL RTU Blocker and Diluent (Vector Laboratories) for 48 hours at room temperature. Primary antibodies (Supplemental Table 1) were diluted (1/200) in RTU Blocker and Diluent for 3 mL total volume added to samples, then incubated for five days at 37°C, then two days at 4°C. Samples were washed in 50 mL PBST at room temperature for five days, exchanging PBST wash buffer each day. Secondary antibodies (Supplemental Table 1) were diluted (1/200) in 5 mL RTU Blocker and Diluent, and samples were protected from light and incubated for five days at 37°C, then two days at 4°C. Samples were washed in 50 mL PBST at room temperature for five days, exchanging PBST wash buffer daily.

Immunolabeled samples were incubated in 20%, 40%, then 63% 2,2’-thiodiethanol (Sigma Aldrich) diluted with PBS at room temperature for two hours each to attain a refractive index of n = 1.45. Samples were then adhered to a glass capillary using cyanoacrylate to mount into the light sheet imaging chamber. Images were obtained with a Zeiss Z.1 Lightsheet microscope using a 20x objective. Images were processed with Arivis Vision4D software (v.3.0.1) and are presented as single optical sections, max intensity projections, and 3D renderings. Quantification of 3D volumes was conducted with Arivis Vision4D in the following manner: image volumes were cropped to isolate structural features of interest, then a supervised intensity-based segmentation was performed using the ‘magic-wand’ feature to isolate channel specific objects, from which volumetric output was derived.

### Data availability statement

The data generated and/or analyzed during the current study are publicly available in the HuBMAP repository. Specifically, the complete CODEX datasets (unprocessed, processed data, along with blank and DAPI channels) with all markers are available on the HuBMAP data portal (https://portal.hubmapconsortium.org/search?entity_type[0]=Dataset).

## Acknowledgements

We thank the families of the organ donors for the gift of tissues. This work was supported by a grant from the NIH NIAID (U54AI142766) to MAA, HSN, TMB, and CHW as well as (P01AI042288) to MAA, The Leona M. and Harry B. Helmsley Charitable Trust (G-2004-03813) to MAA, and JDRF (5-SRA-2018-557-Q-R) to MAA.

## Author Contributions

SC, MJ, JAN, MAB, HH, and CHW performed the experiments and/or analyzed the data. SC, MJ, MAB, JPA, NR, MMS, MY, IK contributed to donor organ acquisition and processing. SC, HSN, CHW, and MAA designed the research. HSN, KO, TMB, CHW, and MAA supervised the research. SC, HSN, and CHW integrated the data and interpreted the results. SC, HSN, CHW, ALP, and MAA wrote the manuscript. All authors discussed the results and edited the manuscript.

## Additional Information

Supplementary information is available for this paper.

## Extended Data Figure and Table Legends

**Extended Data Table 1. Donor demographics, clinical information, and tissue sample identification numbers for SPL and LN examined by IHC, CODEX, and LSFM.**

**Extended Data Figure 1. Immunohistochemical assessment of immune, nervous, endothelial, and stromal markers in human LN and SPL.** CD3ε, CD4, CD8α, CD20, CD21, CD15, Coronin-1A, CD31, CD34, Vimentin, CD107a, CD163, PGP9.5, β3-tubulin, GAP43, ChAT, TH, LYVE1, PDPN and PROX1 were visualized by chromogen-based IHC in human LN (left) and SPL (right) to confirm anticipated staining pattern and determine appropriate antibody dilutions. DAB (brown) indicates a positive signal. Red arrows identify fine neural processes of PGP9.5 within insets of LN and SPL (100 μm).

**Extended Data Figure 2. Immune cell organization within human LN and SPL.** CODEX images of **a,b,c**, LN and **d,e,f**, SPL wide-field (top left, 560 μm) along with high resolution images (30 μm) depicting immune cell distribution throughout tissue parenchyma. **a, d**, CD20 highlights B cells in follicles supported by CD21+, CD35+ FDC meshwork. **b,e**, CD20+ B cells are organized in follicles with admixed CD4+ and CD8+ T cells in the interfollicular space. The periphery of select follicles demonstrates enrichment for CD4+CD45RO+ memory helper T cells, which may represent T follicular helper cells. In the SPL, CD3-CD8α+ cells defining the LCs line the venous sinusoids. **c**, CD15+ neutrophils and CD68+ macrophages are sparsely distributed throughout LN cortex. **f**, In SPL, CD15+ neutrophils are highly abundant both circulating in sinuses and concentrated around follicles throughout the red pulp while CD68+ macrophages are found primarily in the red pulp cords. Donor ID, tissue, and region are defined to the right of each row.

**Extended Data Figure 3. Representative organization of LEC, BEC, and stromal cells in human LN.** Representative CODEX images from five LNs of four independent donors (120 μm). **a**, LYVE1 highlights sinus lymphatics with PROX1 labeling LEC nuclei. LYVE1+ sinusoids, CD4+ T cells admixed with CD8a+ T cells in the perisinusoidal space and Vimentin staining the background stroma (*middle*). CD31 and CD34 identify BECs with SMA accentuating arterioles (*left*). CD163+macrophages in the perisinusoidal space, and CD15+ neutrophils can be seen emerging from the LYVE1+ sinus (*right*). **b-d**, Additional regions from three donor LNs show the distribution of LYVE1 and PDPN expression within perivascular lymphatic sinuses as identified by CD31, CD34, and SMA. **e-g**, Three Additional LN regions from two donors with LYVE1+ sinusoids, CD4+ T cells admixed with CD8a+ T cells in the perisinusoidal space and Vimentin staining the background stroma. **h-j**, Additional LN regions from two donors showing a similar pattern as described in e-g with CD8+CD107a+ T cells and CD163+ macrophages in the perisinusoidal space. Donor ID, tissue, and region are defined to the right of each panel.

**Extended Data Figure 4. Representative organization of LEC, BEC, and stromal cells in human SPL.** Representative CODEX images of five SPL regions from three independent donors (120 μm). **a-c**, LYVE1 highlights splenic lymphatics while SMA and PDPN highlight a trabecula. A CD31+SMA-venules are present and PDPN co-expression is also seen in other smaller SMA+ splenic arterioles. **d-f**, LYVE1+CD8a+ LCs are present with Vimentin highlighting background stroma. Note the distribution of CD4+ T cells around a PALS (unstained, dashed oval, d), early follicle (e), and within the cords of Billroth (f) versus CD8+ T cells. **g-i**, LYVE1 again highlights LCs with CD8a coexpression, and the cords of Billroth show a regular distribution of CD8a+CD107a+ T cells and CD163+ macrophages. Donor IDs and regions are defined to the right of each panel.

**Extended Data Figure 5. Highly multiplexed CODEX images of immune, stromal, and myeloid cells in human LN and SPL along with marker colocalization analysis**. **a**. The LN shows CD20+ B cells organized into follicles with perifollicular CD4+CD3e+CD45RO+ memory T cells (likely T follicular helper) that distribute to nearby CD31+ vasculature. CD3+CD8+ T cells show a somewhat patchy distribution closer to the vasculature and sparing the follicles. HLA-DR+ antigen presenting cells are scattered adjacent to the follicle and vasculature with signal from the CD35+/CD21+ FDCs appearing in the follicles. **b.** The SPL shows a CD31+ arteriole with a white-pulp periarteriolar follicle containing CD20+ B cells and CD21+/CD35+ FDCs. The follicle also contains HLA-DR+ antigen presenting cells, likely dendritic cells. CD8 finely highlights the LCs near which lie CD31+ venules. CD4+CD45RO+ memory T helper cells appear distributed throughout the follicle, rather than the follicle periphery as seen in the LN (a) which implies a difference in follicle age between the two sections. CD3e+CD4-CD8-NK cells are better appreciated in this SPL section than in the LN (a). **c**. LYVE1 highlights LN sinusoidal LECs with focal PDPN expression. PDPN is also identified in the CD21+ follicle. CD31+CD34+ venules and CD31+CD34+SMA+ arterioles are noted. CD68 and CD163 highlight macrophages mostly within the lymphatic sinus and a few CD107a+ interfollicular cytotoxic lymphocytes are noted. Vimentin demonstrates the background stromal architecture (120 μm). **d.** SPL sinuses are demonstrated by CD31, CD34 and LYVE1 with PDPN highlighting a component of an SMA+ arteriole. CD68+CD163+ macrophages and CD107a+ cytotoxic lymphocytes are distributed in the cords of Billroth with a small periarteriolar follicle identified by CD21+ follicular dendritic meshwork. Donor ID and tissue region are defined to the right of each row. **e,f**, Pearson’s R values derived from pair-wise pixel-based colocalization analyses (Coloc2, ImageJ) for multiple protein pairs (N=15 SPL and N=14 LN samples, mean and SD shown in red).

**Extended Data Figure 6. Cell clustering, segmentation, and t-SNE visualization of high-parameter imaging data from human LN.** Parallel analysis of CODEX images from LN (N=4) with **a-d**, donor ID and tissue region defined to the right of each panel. Each panel displays supervised X-shift clustering with identical coloring for each cluster used throughout (*upper left*), heat maps illustrating the protein markers that define each cluster and respective cell counts (*upper middle*), as well as wide-angle and zoomed-in Voronoi diagrams pseudo-colored by cluster assignment (*upper right*). Corresponding t-SNE plots (*lower*) demonstrate global clustering as well as t-SNEs gated for T cells, B cells, neutrophils/macrophages, and endothelial cells. All analyses were performed with the Akoya CODEXMav plugin to NIH Fiji using native algorithm parameters.

**Extended Data Figure 7. Cell clustering, segmentation, and t-SNE visualization of high-parameter imaging data from human SPL.** Parallel analysis of CODEX images from SPL (N=4) with **a-d**, donor ID and tissue region defined to the right of each panel. Each panel displays supervised X-shift clustering with identical coloring for each cluster used throughout (*upper left*), heat maps illustrating the protein markers that define each cluster and respective cell counts (*upper middle*), as well as wide-angle and zoomed-in Voronoi diagrams pseudo-colored by cluster assignment (*upper right*). Corresponding t-SNE plots (*lower*) demonstrate global clustering as well as t-SNEs gated for T cells, B cells, neutrophils/macrophages/littoral cells, and endothelial cells. All analyses were performed with the Akoya CODEXMav plugin to NIH Fiji using native algorithm parameters.

**Extended Data Figure 8. CD31 in SPL highlight endothelium and associated elements entering the *tunica media***. **a**, Photographs of human LN prior to (left) and after (right) tissue clearing using the CLARITY protocol. **b**, Max intensity projection (*left;* 100 μm) and single cross-section (*right*, 50 μm) of a CD31+ blood vessel in spleen. **c**, Max intensity projection of CD31+ splenic blood vessel with DAPI nuclear stain (20 μm). **d**, Cartoon depicting neuronal elements (tyrosine hydroxylase [TH], ChAT, β3-tubulin, and PGP9.5) observed upon the arterial architecture localized to the *tunica externa*, while GAP43 is limited within either the *tunica media* or *tunica intima* (created with BioRender.com; adapted from^56^). **e**, Max intensity projection (*left, center left, and center right*, 50 μm) and cross section (*far right*, 20 μm) of autofluorescence in splenic blood vessels ensheathed by neuronal processes expressing TH demonstrating localization to the *tunica externa*.

**Extended Data Figure 9. LN Lymphatic vessel and nerve fiber networks. a**, LSFM max intensity projections of the extensive β3-tubulin+GAP43+ vessel network (10 μm). **b**, LSFM 3D renderings of LN stained for β3-tubulin and GAP43 (*upper*), β3-tubulin+ vessel with a single GAP43+ nerve fiber overlay (*three lower left panels*, 150 μm), as well as a max intensity projection (75 μm) with inset image of cross-section (50 μm) demonstrating a GAP43+ filament nested within the β3-tubulin+ vessel (*lower right*).

**Extended Data Figure 10. 3D quantification of neuronal markers in vascular segments associated with differential expression and lymphocyte accumulation in human LN and SPL. a**, 3D rendering of blood vessel segments from human LN demonstrating differential expression of GAP43 and β3-tubulin, with the control segment demonstrating GAP43 volume at 55% of total β3-tubulin volume, whereas a target segment demonstrates reduced GAP43 volume at 34% of total β3-tubulin volume. **b**, 3D rendering of blood vessel segment in LN found to have no appreciable β3-tubulin among a GAP43+ vessel. **c**, 3D rendering of blood vessel segments from human SPL demonstrating differential expression of GAP43 and β3-tubulin, with the control segments A and C illustrating GAP43 volume at 41% of total β3-tubulin volume in regions without appreciable Coronin-1A^+^ cells, whereas target segment B demonstrates GAP43 volume at 3% of total β3-tubulin volume in a region displaying significant Coronin-1A^+^lymphocyte accumulation. **d**, An additional vascular segment showing reduced GAP43^+^ volume at 4% of β3-tubulin volume in a region containing Coronin-1A^+^ cell accumulation. Donor IDs are defined to the right of each organ dataset. * Indicates coordination with primary figures.

**Extended Data Figure 11. CODEX data demonstrating lymphocyte (CD20+ B cells and CD45RO+ memory T cells) accumulation around vascular segments in human SPL.** Representative CODEX images of CD31+PDPN+ vascular segments within six CD20+ follicles in a single donor SPL (750 μm). Numbered insets (90 μm) correspond to follicles identified in top left image. Similar findings were observed across donors with bottom row showing follicles from two more donor SPLs (90 μm). Note the distribution of CD45RO+ memory T cells with regard to the vessels and follicle. Donor IDs and tissue CCF region are defined within each representative image.

