## Supplemental Video Legends for "Immune, endothelial and neuronal network map in human lymph node and spleen"

**Supplemental Video 1. 3D LSFM video of dopaminergic neurons in cleared human spleen.** Nerves expressing tyrosine hydroxylase (TH, red) overlayed with autofluorescent vessels shown in cyan (HuBMAP ID 19-01-cc2-a).

**Supplemental Video 2. 3D** **LSFM video of perivascular neurons in cleared human lymph node.** 3D rendering shows overlay of CD31+ vessels (cyan) with nerves expressing choline acetyltransferase (ChAT, yellow) and PGP9.5 (red; HuBMAP ID 20-05-n2-reg01).

**Supplemental Video 3. 3D** **LSFM video of perivascular neurons in cleared human spleen.** 3D rendering shows overlay of CD31+ vessels (cyan) with nerves expressing ChAT (yellow) and PGP9.5 (red; HuBMAP ID 19-01-cc2-a).

**Supplemental Video 4. 3D** **LSFM fly through video of neurons along large and small vessels in cleared human lymph node.** 3D fly through shows overlay of β3-tubulin+ (yellow) and GAP43+ (red) nerves (20-06-n9-reg01).

**Supplemental Video 5. 3D** **LSFM video of neurons and leukocytes in cleared human lymph node.** Nerves expressing β3-Tubulin (yellow) observed along large and small vessels and nerves expressing GAP43 (red) along small vessels only overlayed with Coronin-1A+ leukocytes (cyan; HuBMAP ID 20-06-n9-reg01).

**Supplemental Video 6. 3D** **LSFM video of perivascular neurons in cleared human spleen.** Nerves expressing β3-tubulin (yellow) and GAP43 (red) overlayed with Coronin-1A+ leukocytes (cyan; HuBMAP ID 19-04-cc2-f).
