## Supplemental Table 1 for "Immune, endothelial and neuronal network map in human lymph node and spleen"

**Supplemental Table 1. Antibodies utilized for IHC, CODEX, and LSFM in human spleen and lymph node.**

| **IHC** | | | |  |  |  |  |  |  |
| --- | --- | --- | --- | --- | --- | --- | --- | --- | --- |
| **Target** | **Clone** | **Dilution** | **RRID** |  |  |  |  |  |  |
| CD3e | EP449E | 1/250 | AB_868901 |  |  |  |  |  |  |
| CD4 | EPR6855 | 1/500 | AB_2750883 |  |  |  |  |  |  |
| CD8a | C8/144B | 1/100 | AB_2650657 |  |  |  |  |  |  |
| CD20 | L26 | 1/100 | AB_10734340 |  |  |  |  |  |  |
| CD21 | EP3093 | 1/250 | AB_1523292 |  |  |  |  |  |  |
| CD15 | MMA | 1/200 | AB_397181 |  |  |  |  |  |  |
| Coronin1a | COR1A | 1/500 | AB_2229659 |  |  |  |  |  |  |
| CD31 | EP3095 | 1/250 | AB_2890012 |  |  |  |  |  |  |
| CD34 | QBEnd/10 | 1/1,000 | AB_2861355 |  |  |  |  |  |  |
| Vimentin | RV202 | 1/1,000 | AB_306907 |  |  |  |  |  |  |
| CD107a | H4A3 | 1/100 | AB_1134259 |  |  |  |  |  |  |
| CD163 | EDHu-1 | 1/200 | AB_714951 |  |  |  |  |  |  |
| PGP9.5 | CPCA-UCHL1  polyclonal | 1/500 | AB_2572393 |  |  |  |  |  |  |
| Β_3_-Tubulin | TUJ1 | 1/300 | AB_2313773 |  |  |  |  |  |  |
| GAP43 | CPCA-GAP43  polyclonal | 1/1,000 | AB_2572284 |  |  |  |  |  |  |
| ChAT | GPCA-ChAT polyclonal | 1/100 | AB_2079751 |  |  |  |  |  |  |
| TH | RPCA-TH polyclonal | 1/1,000 | AB_2737417 |  |  |  |  |  |  |
| Lyve1 | EPR21857 | 1/20,000 | AB_2889891 |  |  |  |  |  |  |
| PDPN | NC-08 | 1/100 | AB_1595511 |  |  |  |  |  |  |
| PROX1 | EPR19273 | 1/100 | AB_2868427 |  |  |  |  |  |  |
| **CODEX** | | | | | | | | | |
| **Target** | **Clone** | **Dilution** | **RRID** | **Antibody Cat Number** | **Conjugation Tag** | | **Conjugated In House (Y/N)** | **Cycle #** | **Channel # (CH)** |
| CD31 | EP3095 | 1/200 | AB_1267039 | Akoya 4450017 | AkoyaBX001-AlexaFluor750 | | N | Cycle 2 | CH 2 |
| CD8a | C8/144B | 1/200 | AB_2650657 | Akoya 4250012 | AkoyaBX026-Atto550 | | N | Cycle 2 | CH 3 |
| CD20 | L26 | 1/200 | AB_10734340 | Akoya 4450018 | AkoyaBX007-AlexaFluor750 | | N | Cycle 3 | CH 2 |
| Ki67 | B56 | 1/200 | AB_396287 | Akoya 4250019 | AkoyaBX047-Atto550 | | N | Cycle 3 | CH 3 |
| CD3e | EP449E | 1/200 | AB_764498 | Akoya 4450030 | AkoyaBX045-Cy5 | | N | Cycle 3 | CH 4 |
| SMA | 1A4 | 1/200 | AB_2223019 | ab240654 | AkoyaBX013-AlexaFluor750 | | Y | Cycle 4 | CH 2 |
| Podoplanin | NC-08 | 1/200 | AB_1595616 | Akoya 4250004 | AkoyaBX023-Atto550 | | N | Cycle 4 | CH 3 |
| CD68 | KP1 | 1/200 | AB_11151139 | Akoya 4350019 | AkoyaBX015-Cy5 | | N | Cycle 4 | CH 4 |
| PanCK | AE-1/AE-3 | 1/200 | AB_2616960 | Akoya 4450020 | AkoyaBX019-AlexaFluor750 | | N | Cycle 5 | CH 2 |
| CD21 | EP3093 | 1/200 | AB_1267035 | ab193554 | AkoyaBX032-Atto550 | | Y | Cycle 5 | CH 3 |
| CD4 | EPR6855 | 1/200 | AB_2750883 | Akoya 4350018 | AkoyaBX004-Cy5 | | N | Cycle 5 | CH 4 |
| LYVE1 | EPR21857 | 1/50 | AB_2884014 | ab232935 | AkoyaBX004-AlexaFluor750 | | Y | Cycle 6 | CH 2 |
| CD45RO | UCHL1 | 1/200 | AB_314418 | Akoya 4250023 | AkoyaBX017-Atto550 | | N | Cycle 6 | CH 3 |
| CD11c | 118/A5 | 1/200 | AB_2572997 | Akoya 4350020 | AkoyaBX024-Cy5 | | N | Cycle 6 | CH 4 |
| CD35 | SP197 | 1/100 | AB_2884017 | ab240961 | AkoyaBX016-AlexaFluor750 | | Y | Cycle 7 | CH 2 |
| E-CAD | 4A2C7 | 1/200 | AB_2533118 | Akoya 4250021 | AkoyaBX014-Atto550 | | N | Cycle 7 | CH 3 |
| CD107A | H4A3 | 1/200 | AB_1134260 | Akoya 4350001 | AkoyaBX006-Cy5 | | N | Cycle 7 | CH 4 |
| CD34 | QBEnd/10 | 1/100 | AB_2861355 | NBP2-32932 | AkoyaBX022-AlexaFluor750 | | Y | Cycle 8 | CH 2 |
| CD44 | IM7 | 1/200 | AB_312952 | Akoya 4250002 | AkoyaBX005-Atto550 | | N | Cycle 8 | CH 3 |
| HLADR | EPR3692 | 1/200 | AB_10563656 | Akoya 4450029 | AkoyaBX033-Cy5 | | N | Cycle 8 | CH 4 |
| FoxP3 | 236A/E7 | 1/50 | AB_467556 | 14-4777-82 | AkoyaBX020-Atto550 | | Y | Cycle 9 | CH 3 |
| CD163 | EDHu-1 | 1/200 | AB_714951 | NBI10-40686 | AkoyaBX036-Cy5 | | Y | Cycle 9 | CH 4 |
| COL4 | EPR20966 | 1/200 | AB_2801511 | ab226485 | AkoyaBX029-Atto550 | | Y | Cycle 10 | CH 3 |
| Vimentin | RV202 | 1/400 | AB_306907 | ab8978 | AkoyaBX042-Cy5 | | Y | Cycle 10 | CH 4 |
| CD15 | MMA | 1/200 | AB_397181 | 559045 | AkoyaBX035-Atto550 | | Y | Cycle 11 | CH 3 |
| CD45 | CD45-2B11 | 1/100 | AB_11063696 | 14-9457-82 | AkoyaBX027-Cy5 | | Y | Cycle 11 | CH 4 |
| CD5 | CD5/54/F6 | 1/200 | AB_2884016 | ab213003 | AkoyaBX041-Atto550 | | Y | Cycle 12 | CH 3 |
| CD1c | EPR23189-196 | 1/100 | AB_2884015 | ab270797 | AkoyaBX-30-Cy5 | | Y | Cycle 12 | CH 4 |
| Prox1 | EPR19273 | 1/200 | AB_2868427 | ab236026 | AkoyaBx-002-Atto550 | | Y | Cycle 13 | CH 3 |
| **LSFM** | | | | | | | | | |
| **Primary Antibody** | | | | **Secondary Antibody** | | | | | |
| **Target** | **Clone** | **Dilution** | **RRID** | **Species and Target** | **Wavelength (μm)** | **Dilution** | | **RRID** | |
| CD31 | EP3095 | 1/200 | AB_2890012 | Donkey anti-rabbit IgG | 488 | 1/200 | | AB_2535792 | |
| ChAT | polyclonal | 1/200 | AB_2079751 | Donkey anti-goat IgG | 568 | 1/200 | | AB_2534104 | |
| PGP9.5 | polyclonal | 1/200 | AB_2572393 | Goat anti-chicken IgY | 633 | 1/200 | | AB_2535756 | |
| β3-tubulin | TUJ1 | 1/200 | AB_2313773 | Donkey anti-mouse IgG | 568 | 1/200 | | AB_2534013 | |
| GAP43 | polyclonal | 1/200 | AB_2572284 | Goat anti-chicken IgY | 633 | 1/200 | | AB_2535756 | |
| TH | polyclonal | 1/200 | AB_2737416 | Goat anti-chicken IgY | 633 | 1/200 | | AB_2535756 | |
| Coronin1a | polyclonal | 1/200 | AB_2229659 | Donkey anti-rabbit IgG | 488 | 1/200 | | AB_2535792 | |
