## Extended Data Table 1 for "Immune, endothelial and neuronal network map in human lymph node and spleen"

| HuBMAP Case ID | Donor Age (year) | Donor Sex | Donor Race | BMI | Cause of Death | Organ | Sample ID | Imaging Modality | Figure(s) | HuBMAP Dataset ID | |
| --- | --- | --- | --- | --- | --- | --- | --- | --- | --- | --- | --- |
| 1 | 18 | M | African American | 27.1 | BD | SPL | 19-03-cc2-e28 | CODEX | Ext Data Fig. 5f | HBM724.PJNC.827 | |
| 2 | 0.92 | M | Caucasian | 21.8 | BD | SPL | 19-02-cc2-a28 | CODEX | Ext Data Fig. 5f | HBM968.CJLB.479 | |
| 3 | 18 | M | Caucasian | 25.5 | DCD | SPL | 19-04-cc2-f | LSFM | Fig. 4e-i  Fig. 5d-g  Sup. Video 6  Ext Data Fig. 10c,d | HBM633.BLRX.687 | |
|  |  |  |  |  |  |  | 19-04-cc2-b28 | CODEX | Ext Data Fig. 5f | HBM548.TSMP.663 | |
| 4 | 14 | F | African American | 19.7 | DCD | SPL | 19-01-cc2-a | LSFM | Fig. 3e-h,  Ext Data Fig. 8a-e  Sup. Video 1,3 | HBM363.TMWJ.967 | |
|  |  |  |  |  |  |  | 19-01-cc2-a28 | CODEX | Ext Data Fig. 5f | HBM498.TCSV.345 | |
| 5 | 11 | M | Caucasian | 16.2 | DCD | LN | 20-05-n2-reg01 | LSFM | Fig. 3a-d  Sup. Video 2 | HBM392.QWSF.974 | |
| 6 | 21 | F | African American | NA | BD | LN | 20-06-n1-reg02 | CODEX | Fig. 1a,  Fig. 2a-b  Ext Data Fig. 5e  Ext Data Fig. 6a | HBM866.TGNJ.847 | |
|  |  |  |  |  |  |  | 20-06-n2-reg01 | CODEX | Ext Data Fig. 2a,c  Ext Data Fig. 3b,e,h  Ext Data Fig. 5a,c,e | HBM958.GGVQ.546 | |
|  |  |  |  |  |  |  | 20-06-n3-reg03 | CODEX | Fig. 1b  Ext Data Fig. 3g,i  Ext Data Fig. 5e | HBM355.CZJX.599 | |
|  |  |  |  |  |  |  | 20-06-n9-reg01 | LSFM | Fig. 4a-d  Fig. 5a-c  Sup. Video 4,5  Ext Data Fig. 9a,b  Ext Data Fig. 10a,b | HBM384.XMBW.725 | |
|  |  |  |  |  |  | SPL | 20-06-cc2-c | CODEX | Fig. 1i,m  Fig. 2a-b  Ext Data Fig. 4d,h  Ext Data Fig. 5f  Ext Data Fig. 7a  Ext Data Fig. 11 | HBM556.KSFB.592 | |
|  |  |  |  |  |  |  | 20-06-cc2-b | CODEX | Ext Data Fig. 5f  Ext Data Fig. 11 | HBM568.NGPL.345 | |
| 7 | 20 | M | Caucasian | 28.8 | BD | SPL | 20-07-cc1-a | CODEX | Fig. 2a-b  Ext Data Fig. 4b,i  Ext Data Fig. 5f  Ext Data Fig. 7b | HBM772.XXCD.697 | |
|  |  |  |  |  |  |  | 20-07-cc2-b | CODEX | Fig. 1h,j  Fig. 2a-b  Fig. 5h  Ext Data Fig. 4c  Ext Data Fig. 5b,d,f  Ext Data Fig. 7c  Ext Data Fig. 11 | HBM389.PKHL.936 | |
| 8 | 10 | M | Caucasian | 22.2 | DCD | SPL | 20-08-cc2-b | CODEX | Fig. 1k,n  Fig. 2a-b  Ext Data Fig. 4f  Ext Data Fig. 5f  Ext Data Fig. 7d  Ext Data Fig. 11 | HBM342.FSLD.938 | |
|  |  |  |  |  |  |  | 20-08-cc3-a | CODEX | Fig 1L  Fig 2a-e  Ext Data Fig. 2d-f  Ext Data Fig. 4a,e,g  Ext Data Fig. 5a,f  Ext Data Fig. 7b,d  Ext Data Fig. 11 | HBM772.TKGJ.794 | |
|  |  |  |  |  |  | LN | 20-08-n7-reg03 | CODEX | Fig. 1c  Fig. 2a-b  Ext Data. Fig. 3c,j  Ext Data Fig. 5e  Ext Data Fig. 6b | HBM746.ZTBR.275 | |
|  |  |  |  |  |  |  | 20-08-n9-reg02 | CODEX | Ext Data Fig. 5e | HBM834.ZFVJ.978 | |
|  |  |  |  |  |  |  | 20-08-n10-reg01 | CODEX | Fig. 1e  Fig. 2a-e  Ext Data Fig. 5e | HBM754.WKLP.262 | |
| 9 | 32 | M | Caucasian | 30.2 | BD | LN | 20-09-n1-reg03 | CODEX | Fig. 1d  Fig. 2a-b  Ext Data Fig. 3d,f  Ext Data Fig. 5e  Ext Data Fig. 6c | | HBM957.JXKN.887 |
|  |  |  |  |  |  |  | 20-09-n2-reg02 | CODEX | Fig. 1g  Ext Data Fig. 5e | | HBM747.CQKL.785 |
|  |  |  |  |  |  |  | 20-09-n3-reg01 | CODEX | Ext Data Fig. 5e | | HBM798.MXFT.837 |
| 10 | 47 | M | Caucasian | 28.4 | DCD | LN | 21-10-n3-reg01 | CODEX | Ext Data Fig. 5e | HBM457.JZFF.434 | |
|  |  |  |  |  |  | SPL | 21-10-cc1-a | CODEX | Ext Data Fig. 5f | HBM659.XHFH.996 | |
| 12 | 42 | F | Caucasian | 29.4 | BD | LN | 21-12-n3-reg01 | CODEX | Fig. 2a-b  Ext Data Fig. 5e  Ext Data Fig.6d | HBM723.BZKF.992 | |
|  |  |  |  |  |  | SPL | 21-12-cc3-a | CODEX | Ext Data Fig. 5f | HBM882.DBTR.869 | |
| 13 | 14 | M | Latino | 15.9 | BD | SPL | 21-13-cc2-c | CODEX | Ext Data Fig. 5f | HBM374.LLKS.325 | |
| 14 | 17 | M | Caucasian | 18 | BD | LN | 21-14-n2.5-reg01 | CODEX | Fig. 1f  Ext. Data Fig. 3a  Ext Data Fig. 5e | HBM283.GNGR.785 | |
|  |  |  |  |  |  | SPL | 21-14-cc2-a | CODEX | Ext Data Fig. 5f | HBM543.RSRV.265 | |
| 15 | 20 | M | Caucasian | 23.8 | BD | LN | 21-15-n3-reg01 | CODEX | Ext Data Fig. 5e | HBM483.KDBW.482 | |
|  |  |  |  |  |  |  | 21-15-n3-reg02 | CODEX | Ext Data Fig. 5e | HBM927.KDXB.445 | |
|  |  |  |  |  |  | SPL | 21-15-cc3-b | CODEX | Ext Data Fig. 5f | HBM687.CWVH.758 | |

Abbreviations: M (male), F (female), BMI (body mass index), NA (not available), DCD (donation after circulatory death), BD (brain death), SPL (spleen), LN (lymph node), LSFM (light sheet fluorescence microscopy), CODEX (co-detection by indexing), Ext (extended), Fig (figure), Sup (supplemental).

Sample IDs were assigned according to HuBMAP SOPs. In brief, SPL sample IDs reflect the year the case was acquired, case ID, and common coordinate framework (CCF)-defined region within the tissue. LN sample IDs reflect the year the case was acquired, case ID, node number, and tissue region.
